# Land use impacts poison frog chemical defenses through changes in leaf litter ant communities

**DOI:** 10.1101/745976

**Authors:** Nora A. Moskowitz, Barbara Dorritie, Tammy Fay, Olivia C. Nieves, Charles Vidoudez, Cambridge Rindge and Latin 2017 Biology Class, Masconomet 2017 Biotechnology Class, Eva K. Fischer, Sunia A. Trauger, Luis A. Coloma, David A. Donoso, Lauren A. O’Connell

## Abstract

Much of the world’s biodiversity is held within tropical rainforests, which are increasingly fragmented by agricultural practices. In these threatened landscapes, there are many organisms that acquire chemical defenses from their diet and are therefore intimately connected with their local food webs. Poison frogs (Family Dendrobatidae) are one such example, as they acquire alkaloid-based chemical defenses from their diet of leaf litter ants and mites. It is currently unknown how habitat fragmentation impacts chemical defense across trophic levels, from arthropods to frogs. We examined the chemical defenses and diets of the Diablito poison frog (*Oophaga sylvatica*), and the diversity of their leaf litter ant communities in secondary forest and reclaimed cattle pasture. Notably, this research was performed in collaboration with two high school science classrooms. We found that the leaf litter of forest and pasture frog habitats differed significantly in ant community structure. We also found that forest and pasture frogs differed significantly in diet and alkaloid profiles, where forest frogs contained more of specific alkaloids and ate more ants in both number and volume. Finally, ant species composition of frog diets resembled the surrounding leaf litter, but diets were less variable. This suggests that frogs tend to consume particular ant species within each habitat. To better understand how ants contribute to the alkaloid chemical profiles of frogs, we chemically profiled several ant species and found some alkaloids to be common across many ant species while others are restricted to a few species. At least one alkaloid (**223H**) found in ants from disturbed sites was also found in skins from pasture. Our experiments are the first to link anthropogenic land use changes to dendrobatid poison frog chemical defenses through variation in leaf litter communities, which has implications for conservation management of these threatened amphibians.

## Resumen

Los bosques tropicales, los cuales mantienen la mayor parte de la biodiversidad del planeta, están cada vez más fragmentados por diferentes prácticas agropecuarias. En estos paisajes amenazados por la actividad humana hay organismos que acumulan defensas químicas a partir de su dieta. Por ejemplo, las ranas venenosas (Familia Dendrobatidae) adquieren de su dieta, formada principalmente de hormigas y ácaros de la hojarasca, diferentes alcaloides que pueden ser usados como defensas químicas. Las ranas venenosas están por lo tanto íntimamente conectadas a las redes tróficas locales. Actualmente es desconocido como la fragmentación del hábitat modifica la defensa química a través de cambios en las cadenas tróficas. Por lo tanto, examinamos las defensas químicas y las dietas de la rana diablito (*Oophaga sylvatica*), y las comunidades de hormigas de la hojarasca, en un bosque secundario y en un pastizal cercano. Es destacable que esta investigación fue realizada en colaboración con estudiantes de cursos de ciencias de dos instituciones de educación secundaria. Encontramos que la hojarasca del bosque y del pastizal mantenían comunidades de hormigas con estructuras significativamente distintas. También, encontramos que las dietas y los perfiles de alcaloides en la piel de las ranas en bosque y en pastizal eran significativamente diferentes, las ranas de bosque comían más hormigas (en volumen y número) y acumulaban más alcaloides específicos. Finalmente, la composición de especies de hormigas en las dietas de las ranas fue similar a la de la hojarasca en los alrededores, pero menos variable. Ello sugiere que las ranas tienden a consumir sólo ciertas especies de hormigas en cada hábitat. Para entender mejor cómo las hormigas contribuyen a los perfiles de alcaloides de las ranas, obtuvimos perfiles de los alcaloides presentes en algunas especies de hormigas y encontramos que algunos alcaloides son comunes a muchas especies de hormigas, y otros alcaloides están restringidos a pocas especies. Al menos un alcaloide (**223H**) encontrado en hormigas comunes en áreas disturbadas fue tambien encontrado en la piel de las ranas de los pastizales. Nuestros experimentos son los primeros en vincular los cambios antropogénicos en el uso de suelo con cambios en las defensas químicas de las ranas venenosas a través de cambios en las comunidades de la hojarasca, lo que tiene implicaciones para la conservación de estos anfibios altamente amenazados.

## Introduction

Tropical rainforests boast the highest biodiversity in the world. This biodiversity is supported by intricate food webs whose complexities are only now beginning to be understood [1]. Forest fragmentation due to agriculture and cattle ranching has a large negative impact on biological communities, reducing their diversity and lowering the complexity of local food webs [2–4]. Land use in the tropics is associated with a number of biodiversity declines, including in plants [5], amphibians [6], and arthropods [7]. However, little is known about how changes in land use modifies species interactions across trophic levels [8]. For example, there are many animals in these threatened ecosystems that acquire chemical defenses through their diet, such as poison frogs [9], birds [10], and many arthropods [11]. Chemically defended animals that sequester their toxins are often dietary specialists and thus are dependent on local prey diversity for survival. Studying how land use impacts the trophic interactions mediated by defensive chemicals is important for understanding the factors influencing the stability of local communities and the potential influence of changes in prey item diversity on predator fitness.

Central and South American poison frogs (Family Dendrobatidae) have evolved unique physiological traits that are tightly linked to the trophic structure of rainforest arthropod communities [12]. Poison frogs acquire alkaloid-based chemical defenses from their diet for defense against predators and pathogens [13,14]. Chemical defenses have independently evolved four times within the Dendrobatidae family, coinciding with a dietary specialization on ants and mites in some species [15]. As chemical defenses are environmentally-dependent, there is tremendous variation in the spatial and temporal patterns in the alkaloid profiles across populations within a species [16]. Prior work from our group has established that geographic variation in poison frog chemical defenses reflects chemical diversity of ant and mite communities in their local environment [17]. Thus, it is well established that dendrobatid poison frog chemical defenses are tightly linked to their diet of leaf litter arthropods, but the impact of anthropogenic modification of the environment on these chemical defenses is unknown.

Leaf litter communities comprise a complex set of highly interactive organisms that have major impacts on important ecological functions like nutrient flow [18]. Increasing land use has major impacts on leaf litter communities because it degrades habitat quality [19] and influences trait distribution within communities. For example, a global study examining ant community responses to forest degradation found that both the smallest and largest ant species are lost from communities in degraded habitats [20]. One key aspect of forest degradation is canopy openness [21,22], with most ant groups being lost to habitat disturbance with increasing surface temperature and loss of humidity from the disappearing canopy. Indeed, ants are widely used as bio-indicators of forest quality and restoration success due to their sensitivity to environmental factors correlating with forest health [23]. Although many rainforest animals depend on leaf litter arthropods for nutrients [24, 25], how human-driven changes in leaf litter communities regulate the bottom-up vertebrate community structure and changes in chemical defenses, is not well understood. For example, ant assemblage turnover is predicted to influence alkaloid diversity in the Strawberry poison frog (*Oophaga pumilio*) [26], but there are currently no studies that have quantified poison frog alkaloids and diet and related these measures back to local leaf litter community samples.

In the current study, we utilized a site in Ecuador where the Diablito frog (*Oophaga sylvatica*) spans both secondary forest and reclaimed pasture formerly used for cattle ranching nearly 30 years ago. We sampled frogs in both habitats to ask whether land use impacts frog chemical defenses, diet diversity, and the and species diversity of the surrounding leaf litter. Thus, we determined if differences in frog chemical defenses were due to diet, and how much diet variation could be explained by the composition of leaf litter ant communities. In parallel, we also collected several species of ants to quantify alkaloids typically found in poison frogs to understand how different ant species contribute to poison frog chemical defense. Notably, the dietary arthropod analysis was completed in collaboration with 65 students in two public high schools in the United States as part of a module on ecology. Teachers then participated in follow-up work on habitat quality and frog diet by assisting in the collection of leaf litter ant samples in Ecuador. Overall, our experiments are the first to link anthropogenic land use to modifications in dendrobatid poison frog chemical defenses through changes in leaf litter arthropods.

## Materials and Methods

### Frog sampling

Diablito frogs (*Oophaga sylvatica*) were collected during the day in an area 3.4 km north west of La Florida, Provincia Santo Domingo de los Tsáchilas, Ecuador (−0.250289° −79.027192°) in May and June 2016. Frogs were collected in 100 square meters of secondary rainforest (N=12) and in an adjacent reclaimed cattle pasture (N=9); collection sites are roughly 300 m apart. The secondary forest had dense canopy cover while the reclaimed pasture contained non-native grasses, ferns, and some shrubs and treelets, but no tree cover. In both of these sites, frogs were maintained in enclosures for unrelated research and conservation studies for 2–5 years prior to sampling. The current study only includes female frogs because brain tissues were used for an unrelated study exploring female parental behavior [27], which allowed us to minimize the number of frogs collected and expand the scientific utility of sacrificed individuals. Frogs were captured using plastic cups, anesthetized with a topical application of 20% benzocaine, and euthanized by decapitation. For each individual, the dorsal skin was dissected and stored in methanol in glass vials. The stomach contents were stored in 100% ethanol. Remaining frog tissues were either preserved in 100% ethanol or RNAlater (Thermo Scientific, Waltham, MA, USA) and deposited in the amphibian collection of Centro Jambatu de Investigación y Conservación de Anfibios in Quito, Ecuador. Collections and exportation of specimens were done under permits (Collection permits: 005-15-IC-FAU-DNB/MA and 007-2016-IC-FAU-DNB/MA; CITES export permit 16EC000007/VS) issued by the Ministerio de Ambiente de Ecuador. The Institutional Animal Care and Use Committee of Harvard University approved all animal procedures (Protocol 15-03-239).

### Frog stomach content analysis

Stomach content analyses were completed by high school biology students as part of a laboratory module on ecology, specifically focusing on food webs. Students were trained in their classroom by O’Connell and Moskowitz prior to sample sorting and were supervised the entire time. Students placed frog stomach contents into a petri dish containing phosphate buffered saline (1X PBS) and each student was given 2–10 individual prey items to sort. Each specimen was assigned a unique seven-digit identification number with the first four digits being the frog voucher specimen number and the last three digits being the number assigned to the prey item based on the order in which it was isolated. Each arthropod was photographed with a Lumenera Infinity 2 camera mounted on an Olympus dissecting microscope (SZ40) and stored in 100% ethanol. All prey item photos are available on DataDryad (submission pending).

Diet was quantified by both percent number and percent volume of each arthropod prey type (ants, mites, insect larvae, or “other”) to account for variation in prey size. Adult arthropods that did not fall into discrete dietary categories (adult flies, beetles, etc.) were placed in the “other” category, and all insect larvae were placed in a “larvae” category. Volume was determined by taking length and width measurements from photographs imported into ImageJ (National Institute of Health, Bethesda, Maryland, USA). To verify student measurements, all arthropods were also re-measured by a single individual (Moskowitz). Length was measured from the foremost part of the prey item (including mandibles if applicable) and extended to the rearmost part of the prey item (excluding ovipositors if applicable). Width was measured at the midpoint and excluded appendages. Length and width measurements were then used to calculate the volume of each prey item using the equation for a prolate spheroid: V = (4π/3)*(Length/2)*(Width/2)^2. For spherical mites, volume was calculated using the equation for a sphere: V = (4π/3)*(Radius)^3.

### DNA barcoding of arthropods from stomach contents

DNA of arthropods from frog stomachs (N=407 ants, N=122 mites, N=88 larvae, N=135 “other”) was isolated using the NucleoSpin Tissue kit (Macherey-Nagel, Bethlehem, PA, USA) according to manufacturer’s instructions with a few deviations to make the protocol amenable to a high school classroom schedule as described below. The arthropods were placed individually in a tube containing T1 buffer (from the NucleoSpin Tissue kit), crushed with a pestle, and incubated in Proteinase K solution at 56 °C overnight. The following day, extraction of genomic DNA proceeded according to manufacturer’s instructions and purified genomic DNA was stored at −20°C for one week. We used PCR to amplify a segment of the cytochrome oxidase 1 (CO1 or *cox1*) gene from the mitochondrial DNA, the standard marker for DNA barcoding. CO1 was amplified using the general arthropod primers LCO-1490 (5’-GGTCAACAAATCATAAAGATATTGG) and HCO-2198 (5’-TAAACTTCAGGGTGACCAAAAATCA) from [28]. For all reactions, we used 2 μL of each primer (10 μM) and 25 μL of 2X Phusion High-Fidelity PCR Master Mix with GC Buffer (New England Biolabs, Ipswich, MA, USA) in a total reaction volume of 50 μL. We used a touchdown PCR program to amplify CO1 as follows: 95°C for 5 min; 5 cycles of 95°C for 30 s, 45°C for 30 s with −1°C per cycle, 72°C for 1 min; and 40 rounds of 95°C for 30 s, 40°C for 30 s, and 72°C for 1 min; ending with a single incubation of 72°C for 5 min. PCR reactions were stored at −20°C for one week and then run on a 1% SYBRSafe/agarose gel (Life Technologies). Successful reactions with a single band of the expected size were purified with the E.Z.N.A. Cycle Pure Kit (Omega Bio-Tek, Norcross, GA, USA). Purified PCR products were Sanger sequenced by GENEWIZ Inc. (Cambridge, MA, USA). Arthropod sequences are available on GenBank (MH611377-510 and MN179626-47).

Barcode sequences were imported into Geneious (v 11.0.3) for trimming and alignment of forward and reverse sequencing reactions. We used nucleotide BLAST from the BOLD Identification System (IDS [29], http://www.boldsystems.org/index.php/IDS_OpenIdEngine) to identify our CO1 ant sequences to the genus or species level (Supplementary Excel File). We assigned an order, family, genus or species level annotation based on the results of the BOLD search. We considered results that yielded greater than 96% sequence similarity as sufficient to assign genera or species [30]. For less than 95% BOLD IDS similarity, we assigned specimens to order or family. For certain specimens, results of the IDS search were more taxonomically ambiguous than others. Some specimens only matched to order, rather than a specific family or genus. Additionally, photographs were used to identify each whole ant specimen without barcode sequences to morphospecies level; with successfully barcoded ants used as a reference (Supplementary Excel Table).

### Leaf litter ant survey

Forty leaf litter samples were collected in secondary forest and reclaimed pasture sites in April 2017 at the same sites where frogs had been previously collected (N=20 per habitat type). Each sample consisted of leaf litter removed from randomly placed 1 m^2^ quadrats. Arthropods were extracted using Winkler sacs [31–33], which were loaded and hung for 48 h, during which the arthropods were collected into 70% ethanol. We focused specifically on ants collected from the leaf litter for further analysis. Each specimen was identified to morphospecies using visual inspection against a previously described reference collection of Ecuadorian ants [31–33]. We also obtained samples of 21 species of single ant colonies from the surrounding areas for chemical profiling. When an ant colony was found, we placed 40-100 workers in glass tubes with methanol. For some species, we obtained sufficient samples to run replicates for each colony while for others, we were only able to collect one sample per colony; sample numbers for ant species are in supplementary materials. All ants were visually identified using a reference collection gathered from the site previously [31–33]. Collections (MAE-DNB-CM-2017-0068) and exportation (043-Feb-2018-MEN) of ant specimens were done under permits issued by the Ministerio de Ambiente de Ecuador and Museo de Colecciones Biológicas Gustavo Orcés at Escuela Politécnica Nacional, respectively.

### Alkaloid extraction and quantification

#### Poison frog alkaloids

Frog skin alkaloids were extracted as previously described [17] and stored at −20°C until processing with gas chromatography / mass spectrometry (GC/MS). Samples were analysed on a Quattro micro GC/MS (Waters) fitted with a DB-5 column (30 m, 0.25 mm ID, 0.25 µm film, Agilent). Alkaloid quantification was performed on both forest frogs (N = 10) and pasture frogs (N=7). All raw mass spectrometry data are available on DataDryad (submission pending).

Alkaloids were tentatively identified using the mass spectral data provided in Daly et al [34]. The mass to charge ratio, relative retention times, and relative intensity information was incorporated into a Mass Spectral Transfer File and imported into AMDIS (NIST [35]). This library was used to automatically identify peaks deconvoluted by AMDIS. The identification was weighted by the retention index (calculated from the retention time provided in Daly et al [34]), and the retention index of a few easily identifiable compounds like the Nicotine-d3 internal standard. Manual examination of each candidate toxin’s spectrum was then carried out to improve the accuracy of the candidate identification. Intensities of the model ions for each candidate toxin were extracted from the AMDIS results files and normalized to the mass of skin used for each frog’s alkaloid extraction.

#### Ant alkaloids

Ant alkaloids were extracted as previously described [17] and were analyzed using gas chromatography / mass spectrometry (GC/MS). Samples were analyzed on a Trace 1310 GC coupled with a Q-Exactive Hybrid quadrupole-orbitrap mass spectrometer (ThermoFisher Scientific, Waltham, MA USA). The GC was fitted with a Rxi-5Sil MS column (30 m, 0.25 mm ID, 0.25 µm film, with a 10 m guard column, Restek, Bellefonte, PA, USA). One µl of sample was injected at split 25 in the inlet maintained at 280 °C. The GC program was as follow: 100°C for 2.5 min, then to 280°C at 8°C min^−1^ and held at that final temperature for 5 min. Transfer lines to the MS were kept at 250°C. The MS source was operated in Electron Impact mode, at 300°C. The MS acquired data at 60000 resolution, with an automatic gain control target of 1×10^6^, automatic injection times and a scan range of 33 to 750 m/z.

### Data Analysis

Analyses were conducted using RStudio version 1.1.383 running R version 3.5.2 unless otherwise noted. Boxplots were made with *ggplot2* (version 3.1.0, [36]) and heatmaps were created using the heatmap.2 function in the *gplots* package (version 3.0.1.1).

#### Analysis of arthropods in frog diets

To compare broad arthropod categories in stomach contents across *O. sylvatica* populations, we quantified the relative number and the relative volume of all specimens recovered from the stomach contents, sorted into ants, mites, insect larvae, and *other* arthropods categories (Supplementary Excel File). We visualized assembled arthropod communities using a non-metric multidimensional scaling (NMDS) as available in the R package ‘vegan’ (v 2.5–1, [37]) for both percent volume and percent number of consumed arthropods between habitats. For the NMDS, we calculated Bray-Curtis dissimilarities among samples and plotted the results in two-dimensional plots. Polygons were calculated using the ‘ordihull’ function in ‘vegan’. The purpose of ‘ordihull’ is to create neat, convex outlines to further delineate group separation for visual clarity. A single outline is generated to create a simple polygon that includes all points within an assigned group. To check for differences in the general structure of frog diet categories between habitats, we followed our NMDS with a permutational multivariate analysis of variance (PERMANOVA) on Bray-Curtis dissimilarities. We then checked individual differences between diet categories using a nonparametric Kruskal-Wallis test, given that the data residuals for most diet categories were not normally distributed.

#### Linking ant species in frog stomachs and leaf litter communities

We used an NMDS to explore the relationship between ant species found in frog diets to that of the surrounding leaf litter ant community. We tested for effects of habitat, source and the interaction (habitat x source) with a PERMANOVA on Bray-Curtis dissimilarities.

#### Frog alkaloid analysis

We restricted our alkaloid analysis to fifty reliably identifiable alkaloids that were present in at least five frog specimens (Supplementary Excel File). To confirm alkaloid identification, mass spectra of these candidates of interest were manually inspected and compared to both the Daly database [34] and the NIST14 database [35]. As described above, we visualized samples using a NMDS and tested for differences with a PERMANOVA. We used a nonparametric Kruskal-Wallis to test for differences in alkaloid abundance between forest and pasture frogs. In our analysis of the relative abundances of all 50 alkaloids across groups, p-values were adjusted using the Benjamini-Hochberg procedure to correct for multiple hypothesis testing across all alkaloids.

#### Ant alkaloid analysis

Alkaloids found in the frogs were used as references for a targeted approach to identify alkaloids in ants, meaning that we specifically focused on compounds found in poison frogs rather than all small molecules recovered from the ant alkaloid extraction. For each alkaloid, the molecular ion and main expected fragments were calculated, based on the NIST14 and Daly databases. Retention times for each alkaloid were calculated based on retention indices found in the Daly and NIST14 database. Tracefinder (Thermo) was used to search for the list of alkaloids and to integrate the signal of all potential alkaloids in all the ant data. For each candidate we then selected the sample data with the highest intensity and the spectrum at that retention time was manually evaluated to confirm the identification. We restricted our analysis to alkaloids whose identities could be confirmed and used values normalized by an internal nicotine standard for further analysis. For each species, within-colony replicates were averaged together while between-colony replicates were kept separate.

## Results

As land use continues to alter tropical rainforest ecosystems, we investigated how habitat disturbance influences poison frog chemical defenses and leaf litter arthropod diversity with a special focus on ants. We collected frogs and leaf litter samples in secondary forests and reclaimed pastures to measure poison frog chemical defenses, poison frog diet, and leaf litter arthropods (Figure 1). Overall, we found that land use impacts poison frog chemical defenses through changes in leaf litter species diversity.

**Figure 1.**
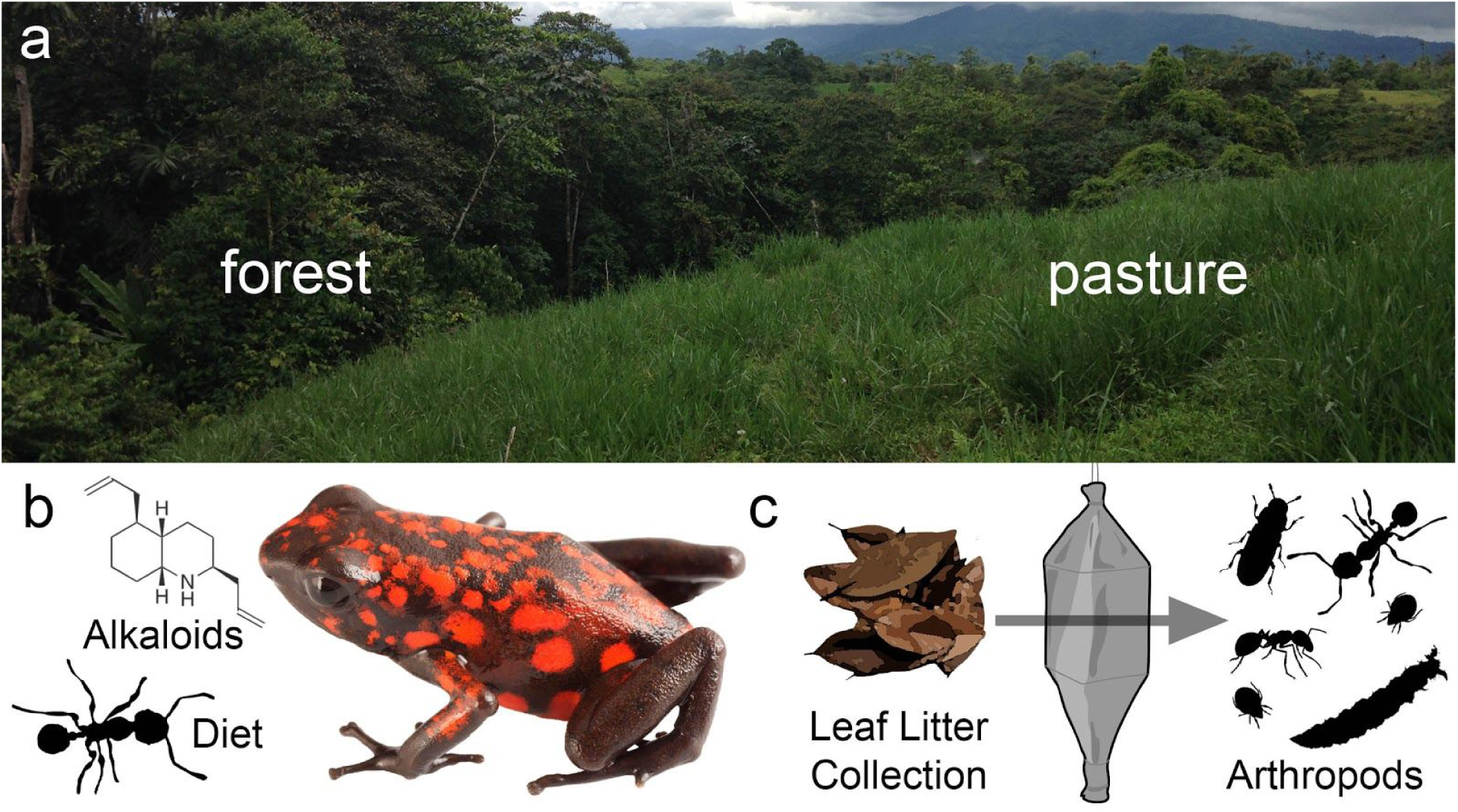
Experimental design to determine the influence of land use across trophic levels. **(a)** Sampling was performed in adjacent enclosures in secondary forest and reclaimed pasture habitats. **(b)** Skin alkaloids and stomach contents were analyzed in frogs from both habitats. **(c)** Leaf litter collections in both habitats were performed using Winkler traps to isolate leaf litter arthropods.

### Diablito frog chemical defenses differ between forest and pasture habitats

Frogs in different habitats showed a significant difference in overall toxin profiles (Figure 2a, NMDS: Stress = 0.097; PERMANOVA: F(1,16) = 3.008, R^2^ = 0.167, p = 0.003). We note that two individuals from the Pasture habitat cluster away from most other samples, but removing these two individuals from the analysis still gives a significant difference between groups (NMDS: Stress = 0.141; PERMANOVA: F(1,14) = 3.787, R^2^ = 0.227, p = 0.003). Among individual alkaloids, 17 differed significantly between frogs from forested and pasture habitats, with 12 surviving false discovery rate correction (Figure 2b). Only three alkaloids were more abundant in pasture frogs (**193D**, **209B**, **223AA**) while all others were more abundant in forest frogs. Decahydroquinolines **221D** (H(1) = 6.223, p = 0.050) and **223**F (H(1) = 7.940, p = 0.035) were both higher in forest frogs. Four 5,8-disubstituted indolizidines differed significantly between groups, with **205A** (H(1) = 9.037, p = 0.025) and **245B** (H(1) = 10.579, p = 0.017) more abundant in forest frogs and **209B** (H(1) = 8.678, p = 0.025) and **223AA** (H(1) = 6.600, p = 0.045) more abundant in pasture frogs. Octohydroquinoline **193D** was more abundant in pasture frogs (H(1) = 8.754, p = 0.025), although this is driven by a few individuals. The tricyclic **245J** (H(1) = 6.617, p = 0.045) and a major isomer (**245J**-like, H(1) = 7.256, p = 0.039) were both more abundant in forest frogs. Finally, 4,6-disubstituted quinolizidine **195C** (H(1) = 10.97, p = 0.017), 2,5 disubstituted pyrrolidine **277D** (H(1) = 6.462, p = 0.046), and the unclassified **253M** (H(1) = 7.559, p = 0.038) were more abundant in forest frogs.

**Figure 2.**
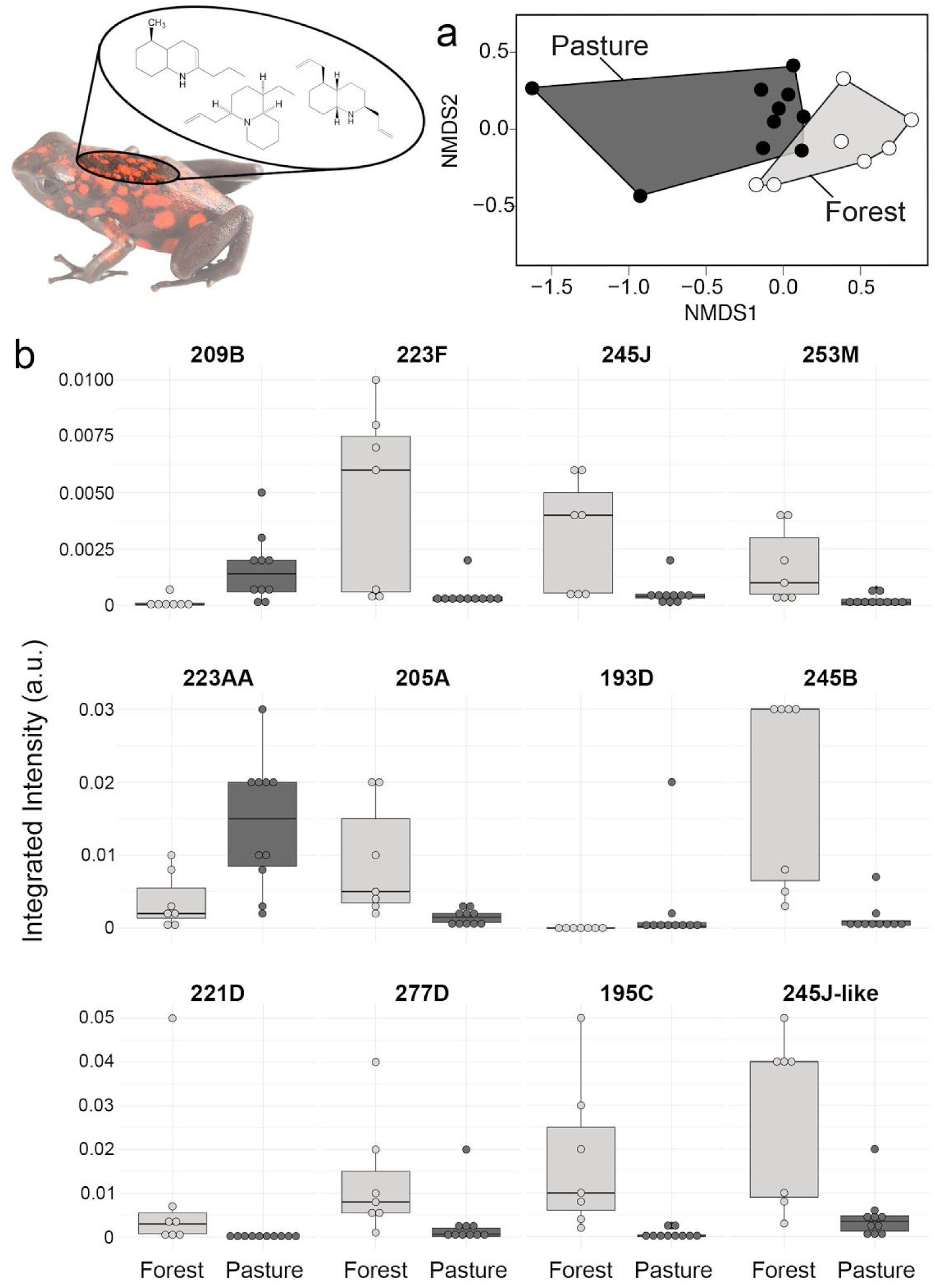
Diablito frog chemical defenses differ between forest and pasture habitats. **(a)** Non-metric multidimensional scaling (NMDS) biplots based on Bray-Curtis dissimilarities show clustering of alkaloid profiles between forest (light grey) and pasture (dark gray) (Stress = 0.10). **(b)** Boxplots show alkaloids that are significantly different in frogs from forest and pasture habitats. Boxplots show the median, first and third quartiles, whiskers (±1.5 interquartile range) and data points (dots).

### Diablito frog diet differs between habitats in number of ants consumed

Forest frogs had more prey items in their stomachs by number and volume compared to pasture frogs (Table 1; number: H(1) = 10.469, p = 0.001; volume: H(1) = 9.778, p = 0.002). Overall differences in relative number of consumed prey categories differed significantly between pasture and forest habitats (Figure 3a, NMDS: Stress = 0.1136; PERMANOVA: F(1,20) = 3.470, R^2^ = 0.1542, p = 0.0258) but relative prey volume did not (Figure 3b, NMDS: Stress = 0.0943; PERMANOVA: F(1,20) = 0.352, R^2^ = 0.0182, p = 0.756). However, relative consumption in both number and volume of individual prey categories varied widely between experimental groups. Forest frog diet comprised of a greater relative number of ants compared to pasture frogs (Figure 3c, Table 1, H(1) = 3.832, p = 0.050), but not a greater relative volume of ants (Figure 3d, Table 1, H(1) = 0.611, p = 0.434), suggesting forest frogs eat more of smaller ant species. Frogs from both habitats did not differ in their percentage consumption of other prey items including mites, larvae or “other” arthropods (Table 1).

**Table 1.** Broad diet characterization of undisturbed and disturbed habitat groups of the Diablito frog, *Oophaga sylvatica*. Diet analyses of different prey types consumed by *Oophaga sylvatica* in forest (N = 12) and pasture (N = 9) habitats. Values show the average across frogs in each group. Significant p-values are in bold. See Supplementary Excel File for full data set. Abbreviations: *H*, H test statistic and p-value from Kruskal-Wallis tests.

|  | Average total number of prey items | % number |  |  |  | Average total volume of prey items (mm<sup>3</sup>) | % volume |  |  |  |
| --- | --- | --- | --- | --- | --- | --- | --- | --- | --- | --- |
|  |  | ants | mites | larvae | other |  | ants | mites | larvae | other |
| **Pasture** | 16.33 | 36.80 | 24.37 | 19.07 | 19.29 | 5.20 | 31.86 | 11.49 | 28.49 | 28.16 |
| **Forest** | 50.42 | 53.71 | 15.51 | 11.57 | 19.76 | 19.29 | 40.16 | 5.61 | 24.86 | 29.37 |
| ***H*** | 10.47 | 3.83 | 3.16 | 0.99 | 0.005 | 9.78 | 0.61 | 2.67 | 0.02 | 0.02 |
| ***p*** | **0.001** | **0.050** | 0.08 | 0.32 | 0.94 | **0.002** | 0.43 | 0.10 | 0.89 | 0.89 |

**Figure 3.**
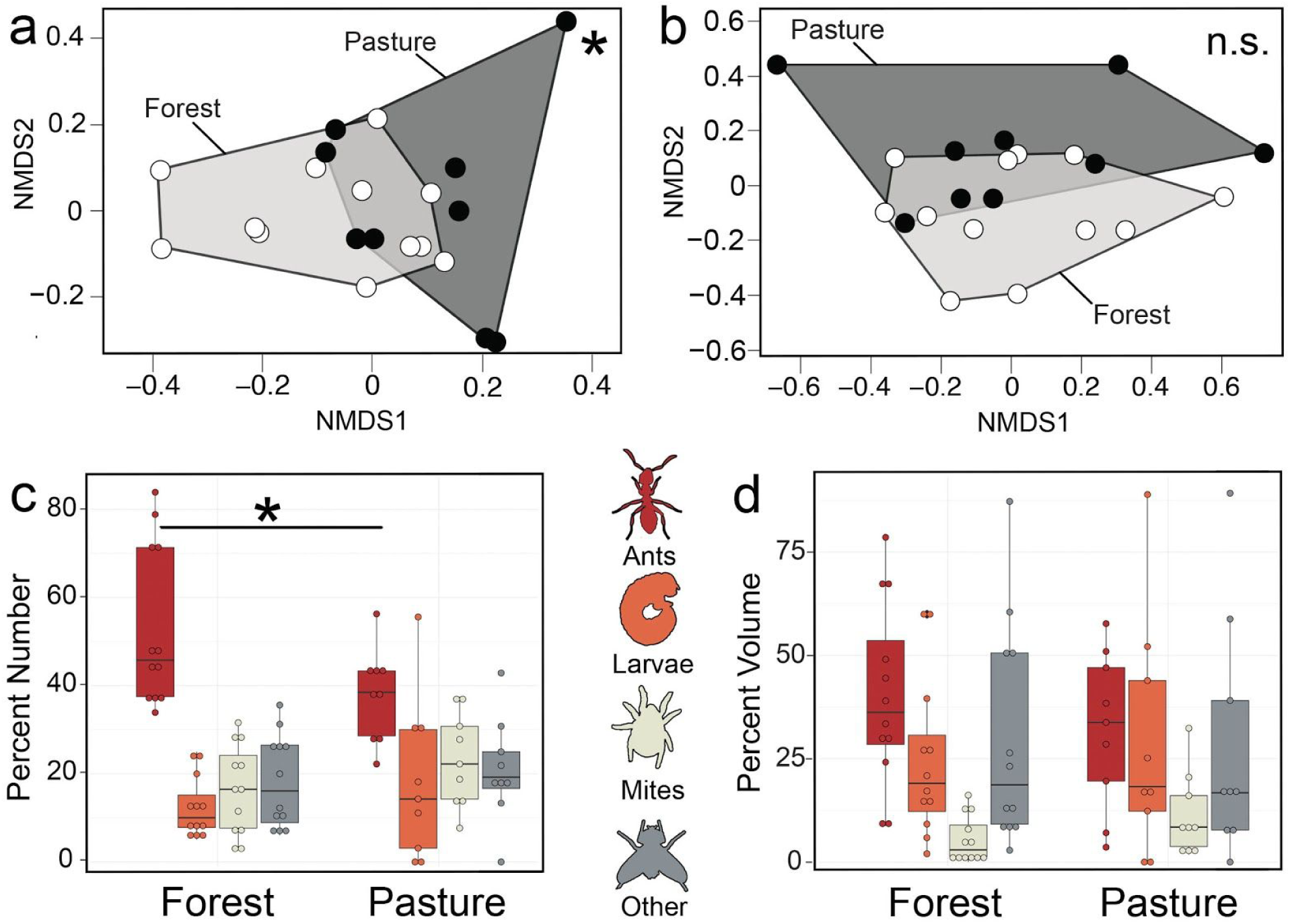
Diablito Frog diet differs between forest and pasture habitats. Non-metric multidimensional scaling (NMDS) biplots based on Bray-Curtis dissimilarities display diet differences between habitats using five prey item categories as input (ants, mites, larvae and “other” prey items) based on (a) percent number (Stress = 0.11) and (b) percent volume (Stress = 0.09) of prey items. Below each NMDS plot is abundance in percent volume (c) and percent number (d) of each prey item category among habitat groups [ants (red), larvae (orange), mites (beige), other (grey)].

### Ant community composition in leaf litter and frog diets

Since many poison frog alkaloids are acquired from ants [17,38], and the number of ants consumed by frogs differed between habitats, we compared ant species diversity in both leaf litter and consumed ants (found in frog stomach contents) in forest and pasture habitats. Total ant abundance recovered from leaf litter spanned 25 ant genera (Supplementary Excel File) and differed between forest and pasture habitats (PERMANOVA: F(1,30) = 3.386, R^2^ =0.101, p < 0.001). Ant composition differed in general between source (frog stomachs vs. leaf litter) (PERMANOVA: F(1,49) = 5.301, R^2^ = 0.086, p = 0.001) (Figure 4a) and between habitats (forest vs. pasture) (PERMANOVA: F(1,49) = 8.136, R^2^ = 0.133, p = 0.001), but only marginally in the interaction (habitat x source) (PERMANOVA: F(1,49) = 1.700, R^2^ = 0.028, p = 0.072) (Figure 4b), which suggests that ants in frog diets are determined only marginally by their availability in surrounding leaf-litter habitats.

**Figure 4.**
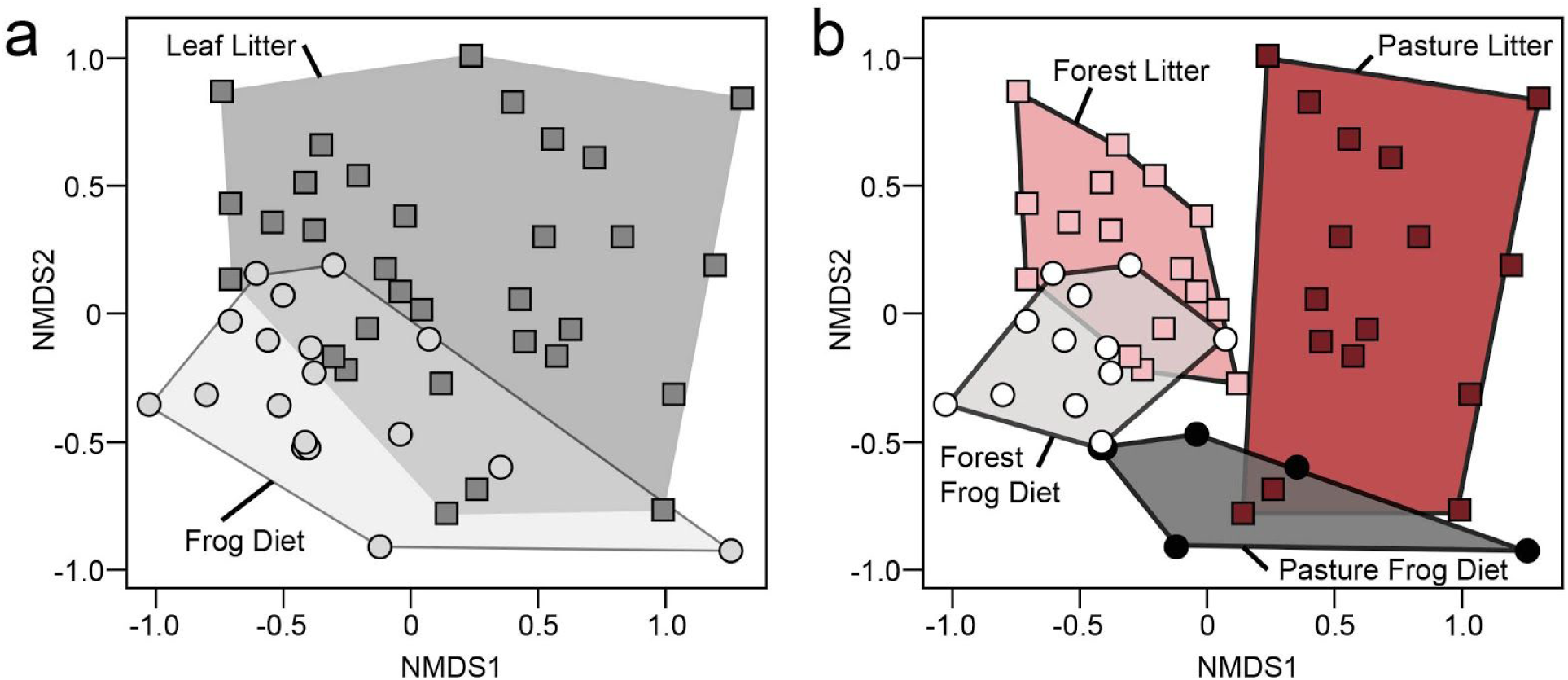
Ant species diversity in the leaf litter and Diablito frog diet differs between forest and pasture habitats. Non-metric multidimensional scaling (NMDS) biplots based on Bray-Curtis distances show ant species diversity differences in **(a)** leaf litter (squares are individual Winkler samples) compared to frog diets (circles are individual frogs) (Stress = 0.19) and **(b)** differences between habitats [pasture litter (dark red), pasture frog diet (dark grey), forest litter (pink), forest frog diet (light grey) (Stress = 0.20).

### Alkaloid diversity across ant species in different habitats

To better understand how variation in ant communities across habitat groups contributes to variation in frog chemical defenses, we profiled alkaloids across 21 ant species, specifically focusing our analysis on alkaloids previously documented in poison frogs (Figure 5a). While the concentrations of many alkaloids were too low to identify with certainty, we confirmed the presence of three alkaloids in our ant dataset: the unclassified alkaloid **191C**, the 2,5-disubstituted pyrrolidine alkaloid **225C**, and a 3,5-disubstituted pyrrolizidine **223H**. The alkaloid **191C** was found in all ants sampled, yet was not found in any frog. The alkaloid **225C** was common across almost all ants in both habitats, although it was not found in *Acropyga* (N=1 colony) and *Strumigenys* (N=1 colony). The alkaloid **223H** was more restricted across ant species, where it was present in most of the *Ectatomma ruidum* and *Linepithema tsachila* samples. Both of these ants have been found in the *O. sylvatica* diet both here and in previous studies [17]. Since both these ant species prefer disturbed or open habitats [39, 40], we predicted that the alkaloid **223H** would differ in abundance between forest and pasture frogs whereas **225C** would not. Indeed, we found that alkaloid **223H** was significantly higher in pasture frogs (Kruskal-Wallis, H(1) = 5.688, p = 0.017) while **225C** was not different between groups (Kruskal-Wallis, H(1) = 3.727, p = 0.054), reflecting the chemical diversity of the ants in their habitats.

**Figure 5.**
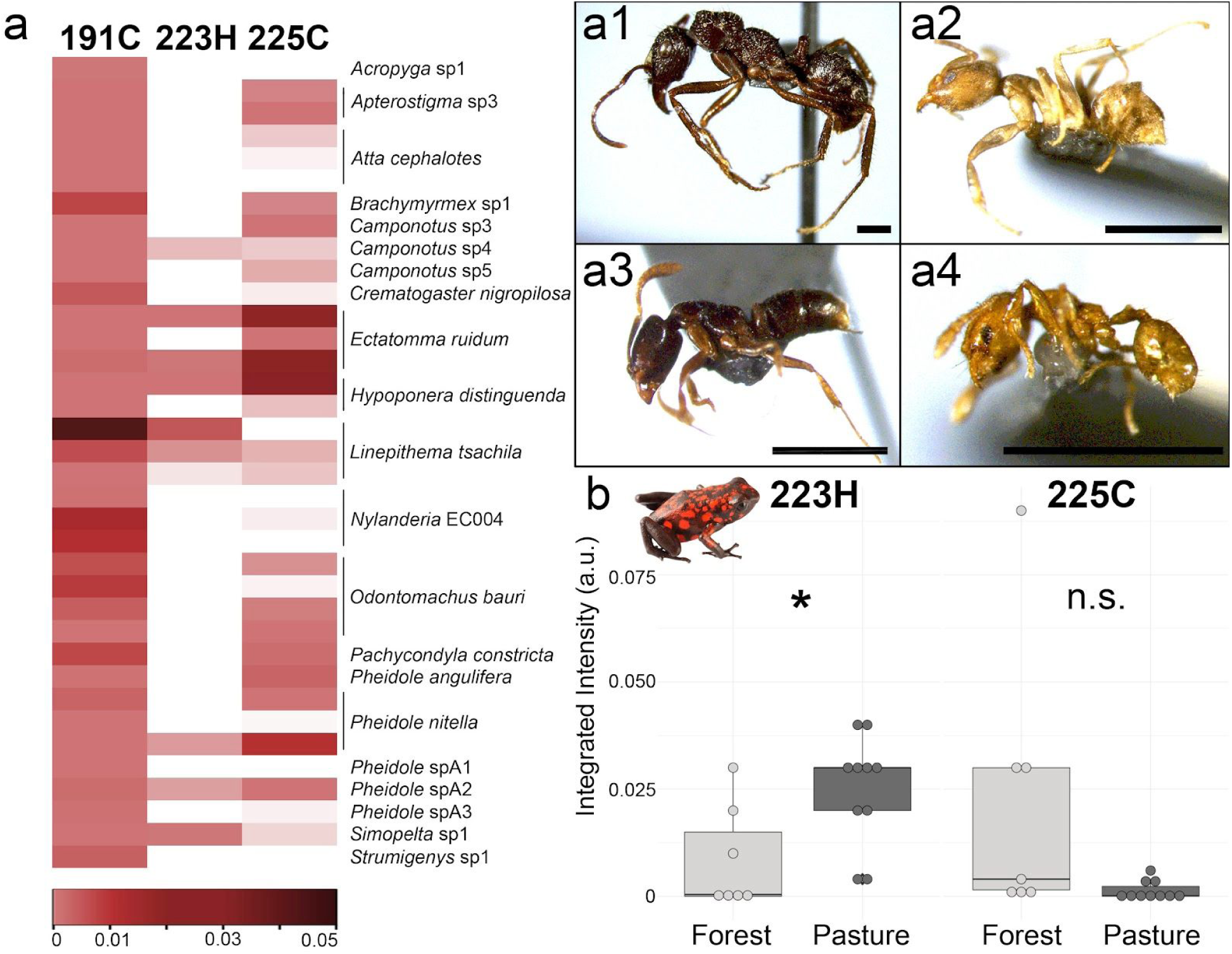
Alkaloid profiles of ants and their abundance in frogs from the same habitat. **(a)** Alkaloids previously described in poison frogs and their abundance across 21 ant species is shown in a heatmap, where alkaloid abundance is plotted in increasing abundance from pink to dark red; white is below the limit of detection. **(a1-a4)** Representative ant species that were positive for **223H** are shown including **(a1)** *Ectatomma ruidum*, **(a2)** *Linepithema tsachila*, **(a3)** *Hypoponera* sp., and **(a4)** *Pheidole nitella*; scale bar 1mm. **(b)** Alkaloids **223H** (right) and **225C** (left) were also detected in frogs. 3,5-disubstituted pyrrolizidine **223H** is significantly more abundant in pasture frogs while there is no difference in 2,5-disubstituted pyrrolidine alkaloid **225C** in frogs from different habitats.

## Discussion

The chemical defenses of poison frogs are acquired from their arthropod diet and are thus susceptible to changing environmental variables. Here, we show that chemical profiles of a dendrobatid poison frog change with anthropogenic land use, likely due to differences in arthropod diet through changes in leaf litter ant communities. We discuss below the influence of land use on poison frog chemical defense sequestration, diet, and leaf litter ecology. Overall our work shows that land use impacts the sequestration of chemical defenses across trophic levels.

### Poison frog diet at the forest-pasture edge

Chemically defended poison frogs are thought to specialize on ants and mites compared to non-defended dendrobatids [41]. At least 50% of their diet is typically composed of these small arthropods, although there is both intra- and inter-specific variation in the proportion of stomach contents made up of ants and mites [17,42,43]. Although previous studies suggest *O. sylvatica* also specialize on ants and mites [17], we found here that other arthropods, such as springtails and insect larvae, also make up a substantial proportion of their diets. Although this is the first study to examine poison frog diet in habitats of differing quality, tree frogs also show dietary variation along successional forest gradients [44], suggesting a trend among tropical amphibians. However, we note the limitation of stomach content analyses as being only a snap-shot in time and more repeated sampling through stomach flushing or fecal barcoding would lend more insight into frog diet across longer periods of time. Regardless, it is clear that variation in habitat quality influences poison frog diet.

Another dimension to dietary differences between habitats is frog foraging behavior. For example, movement may be a contributing factor in diet composition, as the related strawberry poison frog (*O. pumilio*) has been observed to move less in pasture land compared to forest habitats [45]. Indeed, we observed that frogs in the pasture were active for a shorter period of time in the morning compared to the forest frogs. This is reflected in their stomach contents as forest frogs had more arthropods in number and volume than pasture frogs. This difference could be due to either temperature and humidity, as the pasture habitat is hot and dry in the open with little tree cover compared to the closed habitat of the shaded forest that is cooler and more humid. Future studies on the foraging behavior of poison frogs in forest and pasture habitats are needed to disentangle the potential drivers of dietary differences observed here.

### Poison frogs eat a distinct set of leaf litter fauna

Chemically defended poison frogs are thought to have a dietary specialization on ants and mites [41], and to our knowledge, this work is the first in poison frogs to compare stomach contents to leaf litter. We found that forest frogs consumed more ant species than pasture frogs. The few number of ants recovered from pasture frogs’ stomach contents is unexpected because Winkler traps caught a similar number of ant species in pastures and forests (an average of 4.8 and 6.5 ant species, respectively). Furthermore, ant community composition in the leaf litter was significantly different across habitats, suggesting forest and pasture frogs have different ant species available to eat. Moreover, when comparing the diversity of ants consumed by the frogs to those present in leaf litter, we found that frogs consumed a narrow subset of species from the leaf litter ant community. Put together, lower ant diversity in pasture frog stomachs suggests that *O. sylvatica* select particular ants that may be less common in pasture lands. Further tests of these results would benefit from cafeteria assays to test for frog preference.

Most ants consumed by *O. sylvatica* in our study were small and light in coloration (*Solenopsis* and small *Pheidole*). While it is generally unknown if other poison frogs have dietary preferences for particular ant traits, a cafeteria test in mantellid poison frogs suggests they have a size-dependent preference for relatively small prey items [46], a pattern later confirmed by examining stomach contents [47]. These results are troubling, as the smallest ant species (along with the largest ones) are the first to disappear due to habitat modification [20]. However, not all Neotropical frogs choose the smallest ants. For example, non-aposematic *Rhinella alata* frogs in Panama [48] and *Phrynoidis juxtaspera* in Borneo [49] consume larger ants in the leaf litter community. Importantly, in our study, frog preference for certain ant traits did not change with habitat, a pattern also known to occur in Borneo [49]. Therefore, it is unlikely that poison frogs select for ants solely based on their chemical defenses. While there are too few examples to make general conclusions, we note that little is known about top-down control of leaf litter communities and of potential impacts of frog predation on ant communities. Further research would benefit from detailed study of the distribution of relevant alkaloids across ant species within communities and their relations to ant traits. Additionally, for decades the study of frog diets was based on prey identified to broad taxonomic ranks, but we are at the onset of a new era where the advancement of molecular techniques for taxonomic identification of prey items gives us the ability to carefully examine interspecific ant-frog relationships.

### Alkaloids across trophic levels

This work is the first to our knowledge to examine how land use influences chemical defenses in a dendrobatid frog. We found that forest frogs have more specific kinds of alkaloids than pasture frogs, although there are a few alkaloids that are more abundant in pasture frogs. There are two conflicting reports regarding habitat quality and chemical defenses in Malagasy mantellid frogs, an independent evolutionary origin of acquired chemical defenses in amphibians [50]. One study reports increased alkaloid diversity and quantity in mantellids from undisturbed forest habitats compared to disturbed habitats [51]. The other study reports that mantellid frogs in disturbed habitats had greater alkaloid diversity [52], although this trend was only observed in one of three sites. These studies also differed in the method of alkaloid sampling (whole skin versus non-invasive sampling) making direct comparisons difficult. While more research at forest-pasture edges is necessary to make general conclusions about habitat disturbance and chemical defenses, these data clearly indicate that environmental changes can impact poison frog alkaloid profiles. Future work should also consider potential consequences for poison frog fitness to determine if conservation strategies should be developed to address these anthropogenic effects.

Most alkaloids sequestered by poison frogs have an ant or oribatid mite origin [9]. Although a broad analysis of ant alkaloids has never been performed across many tropical ant species, formicine and myrmicine ants carry some alkaloids sequestered by dendrobatid frogs [38,53]. It is possible that the greater abundance and diversity of ants consumed by forest frogs compared to pasture frogs contributes to the differences in chemical defenses observed in these forest frogs, although this is difficult to test with the present data set. We reliably found three poison frog alkaloids in some of the 21 species of ants that were randomly surveyed within the frog collection sites. The unclassified alkaloid **191C** was found in all ant species, but was not detected in any frogs. To our knowledge, this is the first report of **191C** in ants. It is therefore surprising that *O. sylvatica* did not carry this alkaloid given all ants sampled contained detectable levels. Although poison frogs do uptake many different alkaloids, this process is not ubiquitous, where feeding experiments have shown some specificity in alkaloid uptake [54]. *O. sylvatica* may be unable to sequester **191C**, but controlled feeding experiments in the laboratory are needed to test this. We also detected **223H** and **225C** in the ants sampled here and these alkaloids have been reported in ants previously [34]. Alkaloid **225C** was present across many ants while **223H** was restricted to fewer species. Moreover, the habitat preferences and alkaloid diversity of these ants predicted alkaloid abundance observed in frogs from different habitats. Finally, we note that there is substantial variation in alkaloid presence across ant colonies within the same species. Since many alkaloids are synthesized by microbes and plants [55,56], it is possible that ants also acquire these alkaloids, or their precursors, from environmental sources like diet, which is currently unknown and should be an area of future research effort.

### Future directions in poison frog conservation

Many open questions remain from this study that could influence future conservation efforts of poison frogs and their tropical forest habitat. First, it is currently unknown how forest-pasture edge effects influence *O. sylvatica* distribution. Amphibian species seem to vary in whether edge effects influence their spatial distributions [57], and future population surveys across many sites are needed to better understand poison frog distributions around the forest edge. Second, given the results presented here, long term monitoring of poison frog chemical defenses in regions with human land use is needed. Although poison frog chemical defenses are environmentally derived, their bright aposematic coloration is genetically encoded. Thus, loss of alkaloid-sources from the diet would render poison frogs undefended but still brightly colored. This bright coloration coupled with no chemical defenses may put the frogs at risk of higher predation from random sampling by predators and subsequent relaxed associative learning. More research on poison frog predation in the context of aposematism is needed to make accurate conservation predictions. As anthropogenic change continues to impact the habitats of many chemically defended animals, a priority for future basic science and applied conservation research should focus on how chemical defense is impacted by land use.

### Data Accessibility

All data is available in the Supplementary Excel File. Arthropod photos and mass spectrometry data are available on DataDryad (submission pending).

## Acknowledgements

We would like to thank Lola Guarderas for logistical support in Ecuador, Jules Wyman for the ant photographs, and Callie Chappell and Daniel Friedman for comments on earlier versions of this manuscript.

## Author Contributions

LAO, LAC and DAD designed the research; EKF and NAM collected frog samples in the Ecuador; DAD, BD and TF collected leaf litter ant specimens; BD and TF’s high school biology students performed the stomach content measurements and DNA barcoding under the guidance of NAM and LAO. LAC maintains the enclosures in forested and pasture habitats; DAD performed morphological identification of ant specimens and conducted the leaf litter diversity analysis; CV and SAT performed the small molecule mass spectrometry and analyses; NAM compiled the data files, contributed to the diet analyses, and managed the project; DAD and LAO analyzed the data and wrote the manuscript with contributions from all authors.

Cambridge Rindge and Latin 2017 Biology Class (Teacher: Barbara Dorritie): Giavanna Benitez, Jack Deford, Rory Fitzpatrick, Lucas Guzman-Finn, Clara Iffland, Na-Jae Josephs, Maximillian Kaufman, Guillermo Lopez, Tyler Marcus, Keilah Michel, Aaron Pen, Abigail Powers-Lowery, Pablo Reina-Gonzalez, Aidan Richards, Abdelaziz Rifai, Smarika Suwal, William Telingator, Kenya Wade, Julian Warburton, Abenezer Zewede, Margaret Allen, Fosca Bechthold, Tal Ben-Anat, Lucy Bent, Ian Chester, Ruth Desta, Mekinsa Frith, Karla Goss, Frederick Gould, Jeynaba Jamanka, Nadaija Lauture, Katie Melendez, Kyle Mercado, Ivan Munzert, Keanan Ng, Marina Pineda-Shokooh, Danielle Reeves, Natalia Ruiz, Dagmawi Wassie, Evan Wilcox, Elaina Wolfson.

Masconomet 2017 Biotechnology Class (Teacher: Tammy Fay): Ariana Afshar, Benjamin Anderson, Sherri Blais, Christian Bovest, Terrence Bovest, Nicholas Celso, Jack Connors, Emily Constan, Christina Curreri, Benjamin Demers, Gwen Ellis, Mary Erb, Marcos Esperon, Mia Farris-Kindregan, Nicholas Messina, Nicholas Migneault, Anthony Nazarian, Olivia Nieves, Lily O’Grady, Aubrey O’Keefe, Thomas O’Keefe, Thomas Pappalardo, Brock Reardon, Luca Shanley.

